# Sugarcane Nitrogen and Irrigation Level Prediction Based on UAV-Captured Multispectral Images at the Elongation Stage

**DOI:** 10.1101/2020.12.18.423409

**Authors:** Xiuhua Li, Yuxuan Ba, Shimin Zhang, Mengling Nong, Muqing Zhang, Ce Wang

## Abstract

**Introduction:** Sugarcane is the main industrial crop for sugar production; its growth status is closely related to fertilizer, water, and light input. Unmanned aerial vehicle (UAV)-based multispectral imagery is widely used for high-throughput phenotyping because it can rapidly predict crop vigor. This paper mainly studied the potential of multispectral images obtained by low-altitude UAV systems in predicting canopy nitrogen (N) content and irrigation level for sugarcane.

**Methods:** An experiment was carried out on sugarcane fields with three irrigation levels and five nitrogen levels. A multispectral image at a height of 40 m was acquired during the elongation stage, and the canopy nitrogen content was determined as the ground truth. N prediction models, including partial least square (PLS), backpropagation neural network (BPNN), and extreme learning machine (ELM) models, were established based on different variables. A support vector machine (SVM) model was used to recognize the irrigation level.

**Results:** The PLS model based on band reflectance and five vegetation indices had better accuracy (R=0.7693, root mean square error (RMSE)=0.1109) than the BPNN and ELM models. Some spectral information from the multispectral image had obviously different features among the different irrigation levels, and the SVM algorithm was used for irrigation level classification. The classification accuracy reached 77.8%.

**Conclusion:** Low-altitude multispectral images could provide effective information for N prediction and water irrigation level recognition.

## 0 Introduction

As the most important sugar crop, sugarcane is mainly grown in tropical and subtropical areas and provides approximately 80% of the world’s sugar. The growth of sugarcane is closely related to fertilizer, water, and radiation intensity. Evaluating the growth situation of sugarcane in a timely manner and adjusting the field management strategy accordingly is of great significance to the yield and quality of sugarcane. In recent years, remote sensing of different scales with spectral images has been considered an effective high-throughput phenotyping solution and has been widely used to predict the growth potential or yield of different crops.

Most of the earliest spectral images were obtained from satellite-based acquisition platforms, such as Sentinel [1], Gaofen (GF) [2], Landsat [3,4], GeoEye [5], and QuickBird [6], normally provided by governments or commercial companies. Their main agricultural applications include land cover and land use monitoring, vegetation classification, yield prediction, and growth prediction. However, although satellite remote sensing can obtain crop population information on a large scale, due to the limitations of a low spatial resolution and fixed revisit cycles, it has formidable deficiencies for small-scale applications, which usually need more subtle information or a higher temporal resolution for crop growth monitoring [7].

In the early 21^st^ century, many well-funded research institutions began to use aviation aircraft to carry multispectral or even hyperspectral cameras at an altitude of several to tens of kilometers to acquire spectral images and conduct many agricultural research and applications [8,9]. However, this method was difficult to popularize due to its relatively high cost.

With the rapid development of unmanned aerial vehicle (UAV) technology and the increasing availability and decreasing cost of spectral image sensors, there are increasingly favorable opportunities for capturing high-spatial-resolution and high-spectral-resolution image data. UAV-based remote sensing systems have a much higher ground resolution (centimeter-level), which means they are more sensitive to spatially heterogeneous information due to their lower flying altitudes. Over the past 10 years, UAVs, especially drones, have been rapidly accepted and popularized to acquire reliable field crop growth information, weather permitting. This technology can compensate for traditional crop monitoring equipment problems, such as small scope and great difficulty, and has good application value. Multispectral imaging technology has been used for crop quality detection due to its advantages of fewer spectral bands, short information acquisition time, simple data structure, etc. [10].

Many previous studies have shown that rice 11, corn [12], barley [13], and wheat [14] biomass could be reliably predicted by using multispectral and RGB images obtained by drones. Canopy spectral information was effective for estimating the grain protein content of winter wheat (R^2^=0.7939) [15]. Ballester et al. evaluated the effectiveness of a set of spectral indicators obtained from UAVs in tracking temporal and spatial variations in nitrogen (N) states and predicting commercial cotton field yields [16]. Hyperspectral techniques have been used to detect crop diseases [17] and estimate N concentrations in canopy leaves [18].

Similar studies have also been conducted in recent years for sugarcane [19]. Some researchers used satellite imagery [20,21]21 and UAV hyperspectral imagery [22,23] to predict sugarcane biomass and achieved varying degrees of success. However, no reports have been published on N and irrigation level prediction for sugarcane based on UAV imagery.

Partial least squares (PLS), backpropagation neural networks (BPNNs), and extreme learning machines (ELMs) have been widely used in nutrient predictions. For example, Mahanti et al. [24] used PLS to predict the nitrate content of spinach, and Li et al. [25] used PLS to establish 12 models of fruits and seeds for rapid analysis and quality assessment. Kira et al. [26] established a model for estimating the chlorophyll and carotenoid contents of three tree varieties based on a BPNN. Chen et al. [27] constructed a BPNN model to invert rice pigment content with several spectral parameters as input. Pal et al. [28] proposed the ELM algorithm to classify land covers with multispectral and hyperspectral data; it achieved a better classification accuracy than models established with BPNN and support vector machine (SVM), with far less computational complexity.

Therefore, in this study, high-resolution multispectral images of an experimental sugarcane field were obtained by a low-altitude UAV, and prediction methods of the sugarcane canopy N content were compared and evaluated by PLS, BPNN, and ELM models. Additionally, different irrigation levels were preliminarily identified and classified by using multispectral images.

## 1 Material and methods

### 1.1 Experimental design and image acquisition

The sugarcane experimental field was located in Nanning, Guangxi Autonomous Region, China (latitude 22.84° N, longitude 108.33° E). Two irrigation treatments and four fertilization treatments were applied in the field. Urea, calcium magnesium phosphate, and potassium chloride were chosen as the N, phosphorus (P), and potassium (K) fertilizers, respectively. Eight plots with different irrigations and fertilizers and four blank plots (denoted by BL, without fertilizer and drip irrigation) were included in the field. Concrete partitions at a depth of 1.2 m were built between each plot to prevent water and fertilizer infiltration. The two irrigation treatments included 180 m^3^/ha (denoted by W0.6) and 300 m^3^/ha (denoted by W1.0). The four fertilizer treatments were as follows: F1.0: 250 kg/ha of N, 150 kg/ha of P_2_O_5_, 200 kg/ha of K_2_O; F0.9: 10% reduction in the amount of F1.0; F1.1: 10% increase over F1.0; F1.2: 20% increase over F1.0. Water and fertilizer were applied via drip irrigation pipes. The same amount of micronutrient fertilizer was applied to all the plots except the blank plots. The eight plots were the same size of 6.6 m×5.6 m, with the different treatments denoted by W0.6F0.9, W0.6F1.0, W0.6F1.1, W0.6F1.2, W1.0F0.9, W1.0F1.0, W1.0F1.1 and W1.0F1.2. The seed canes were planted on March 24, 2018, seedling fertilizer was applied on May 11 (48 days after planting), and tillering fertilizer was applied on June 29 (97 days after planting).

The multispectral images were captured at noon on July 11, 2018 (109 days after planting), in the elongation stage. The image acquisition system was mainly composed of a Phantom 4 Pro drone (DJI, China) and a RedEdge-MX multispectral image sensor (MicaSense, USA), as shown in Fig 1A and 1B, respectively. The RedEdge-MX imager has five spectral bands at 475 nm, 560 nm, 668 nm, 717 nm, and 840 nm and is equipped with a light intensity sensor and a fixed reflectivity correction panel (Group VIII, USA) for radiation correction. The optical intensity sensor could correct the change in the influence of external light on the spectral image, and the fixed reflectance correction panel could perform reflectance transforms. The weather was sunny and windless. The drone flew at an altitude of 40 m, with 85% forward and 85% side overlap. The time interval of image acquisition was 2 s, and the ground sample distance (GSD) was 2.667 cm. In total, 260 images were collected and mosaiced in Pix4Dmapper (Pix4D, Switzerland). The false-color mosaic image is shown in Fig 2. Three to four first leaves were collected from adjacent stalks as one sample, and three random samples from each plot, 36 samples in total, were collected for N determination.

**Fig 1.**
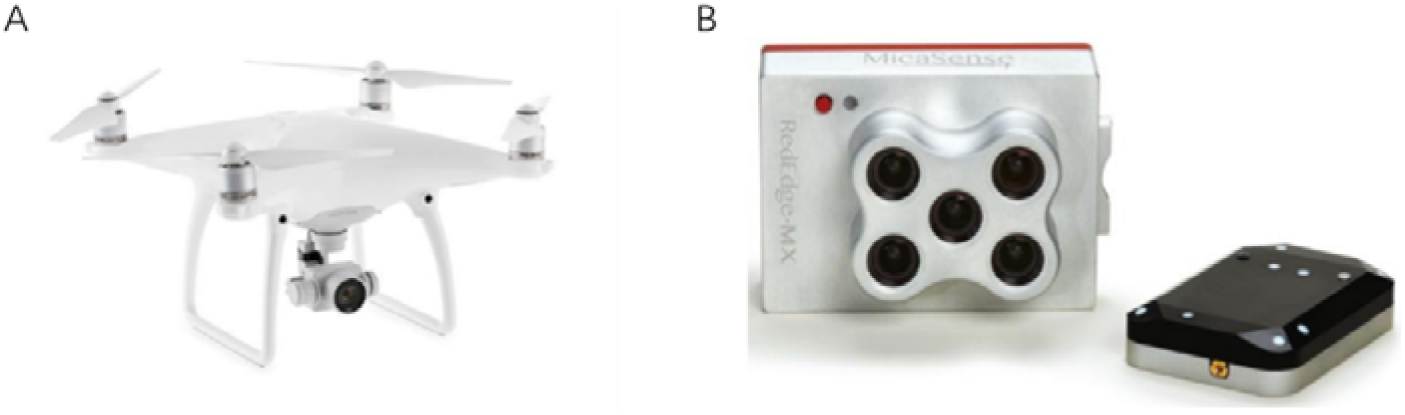
The image acquisition system consisted of a drone (A) and a multispectral sensor (B).

**Fig 2.**
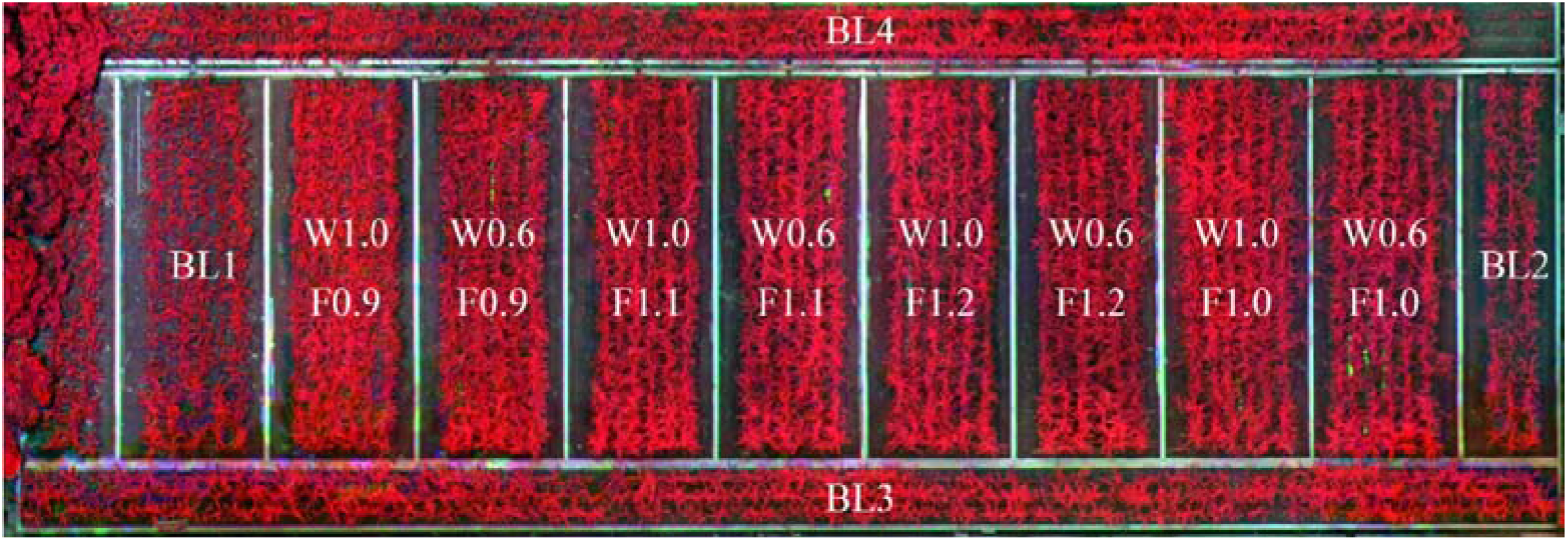
The false-color mosaic image of the sugarcane field and the plot distribution. BL means blank, W means water irrigation, F means fertilizer application.

### 1.2 Multispectral image preprocessing

The mosaic image was mainly processed in ENVI software (L3Harris Geospatial, USA). Radiation calibration was first implemented using the radiometric correction module. Atmospheric correction was conducted using the fast line-of-sight atmospheric analysis of hypercubes (FLAASH) tool to eliminate the radiation errors caused by atmospheric scattering and other factors to a certain extent.

The normalized difference vegetation index (NDVI) image was calculated by simple band math. The decision tree (DT) algorithm was used to extract the sugarcane canopy from the soil, weeds, shadow, and other interfering factors. Fig 3 shows the extraction result, and the white dots in the figure represent the sample locations.

**Fig 3.**
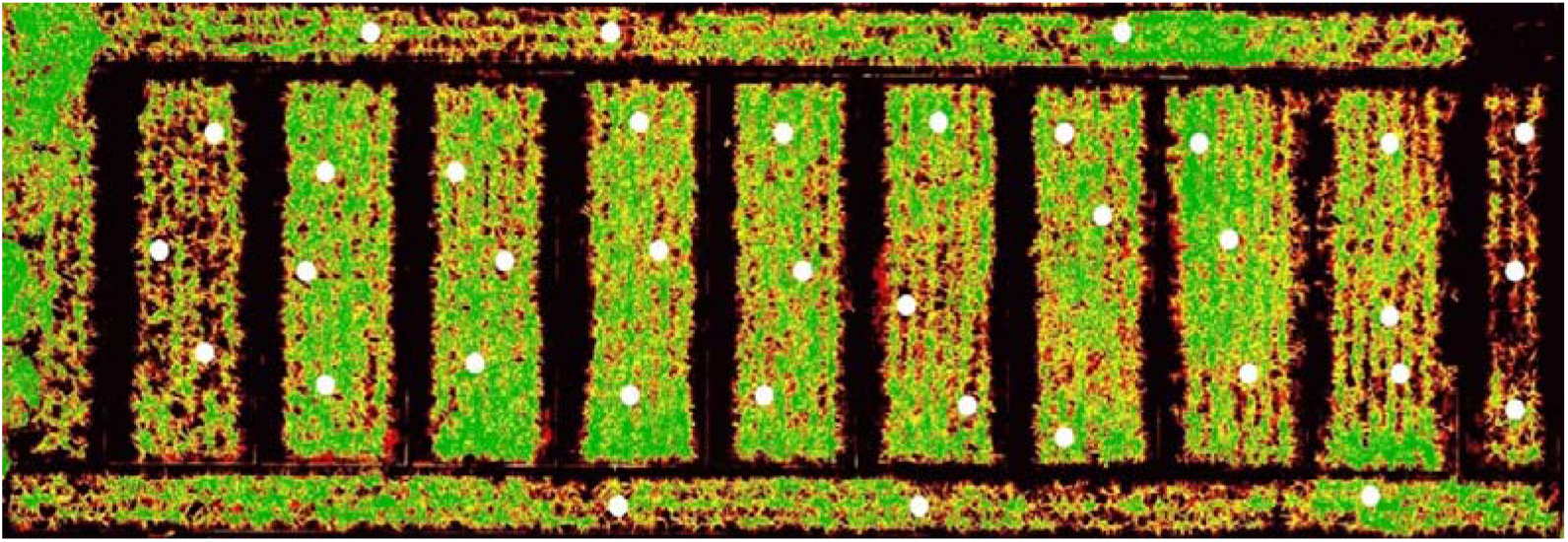
Sugarcane canopy extraction result based on the NDVI. The white dots represent the sample locations.

### 1.3 Modeling algorithms

PLS is an algorithm that combines the advantages of multiple linear regression and canonical correlation analysis and principal component analysis, commonly used when the model input variables are large in number and have multiple correlations. The model is settled at the minimum values of the sum of square error. The backpropagation neural network (BPNN), namely, the multilayer feedforward neural network based on the error inverse propagation algorithm, could theoretically approximate any nonlinear continuous function with arbitrary precision, which had a strong fitting ability for nonlinear functions and features of self-learning and self-adaptation. The neural network model had a strong anti-interference ability and may be suitable for complex field environments. The ELM algorithm allows the random generation of the weights and threshold between the input layer and hidden layers; users need to specify only the number of hidden layer neurons in the whole training process. Compared with traditional classification algorithms, ELM has a fast learning speed and strong generalization capability.

## 2 Results and discussion

### 2.1 Nitrogen content prediction based on multispectral images

The average spectral or VI value of an area of 0.1 m by 0.1 m, with ground sample location as its center, was calculated as the counterpart to the ground sample to reduce random error.

#### 2.1.1 Modeling directly with the five spectral bands

The five multispectral image bands were first directly taken as the input variables, and the PLS, BPNN, and ELM algorithms were used to build canopy N content prediction models. Threefold cross-validation was used to evaluate accuracy. The PLS model had a higher R value and lower root mean square error (RMSE) and standard prediction error (SEP) value in the validation set than the ELM and BPNN models, indicating that it has a relatively higher accuracy. The ELM and BPNN models had R values of 1.000 and very low RMSEs, showing dramatically better results than the validation set, indicating a high probability of overfitting (Table 1).

**Table 1.** PLS, ELM, and BPNN modeling results based on the five spectral bands.

| Modeling algorithms | Training |  |  | Validation |  |  |
| --- | --- | --- | --- | --- | --- | --- |
|  | R | RMSE | SEP | R | RMSE | SEP |
| PLS | 0.9092 | 0.0653 | 0.0667 | 0.6986 | 0.1166 | 0.1209 |
| ELM | 1.0000 | 0.0006 | 0.0026 | 0.5698 | 0.1455 | 0.1497 |
| BPNN | 1.0000 | 0.0004 | 0.0034 | 0.5706 | 0.1590 | 0.1523 |

#### 2.1.2 Modeling with vegetation indices

Vegetation indices (VIs) have been widely used to qualitatively or quantitatively evaluate vegetation cover varieties and vigor. The NDVI is the most commonly used VI, and it is also one of the important parameters closely related to crop chlorophyll and N contents.

To compare the effects of different VIs on predicting canopy nutrient content and find the optimal VI or a combination of VIs, 10 commonly used VIs (as shown in Table 2) were selected for further study.

**Table 2.**
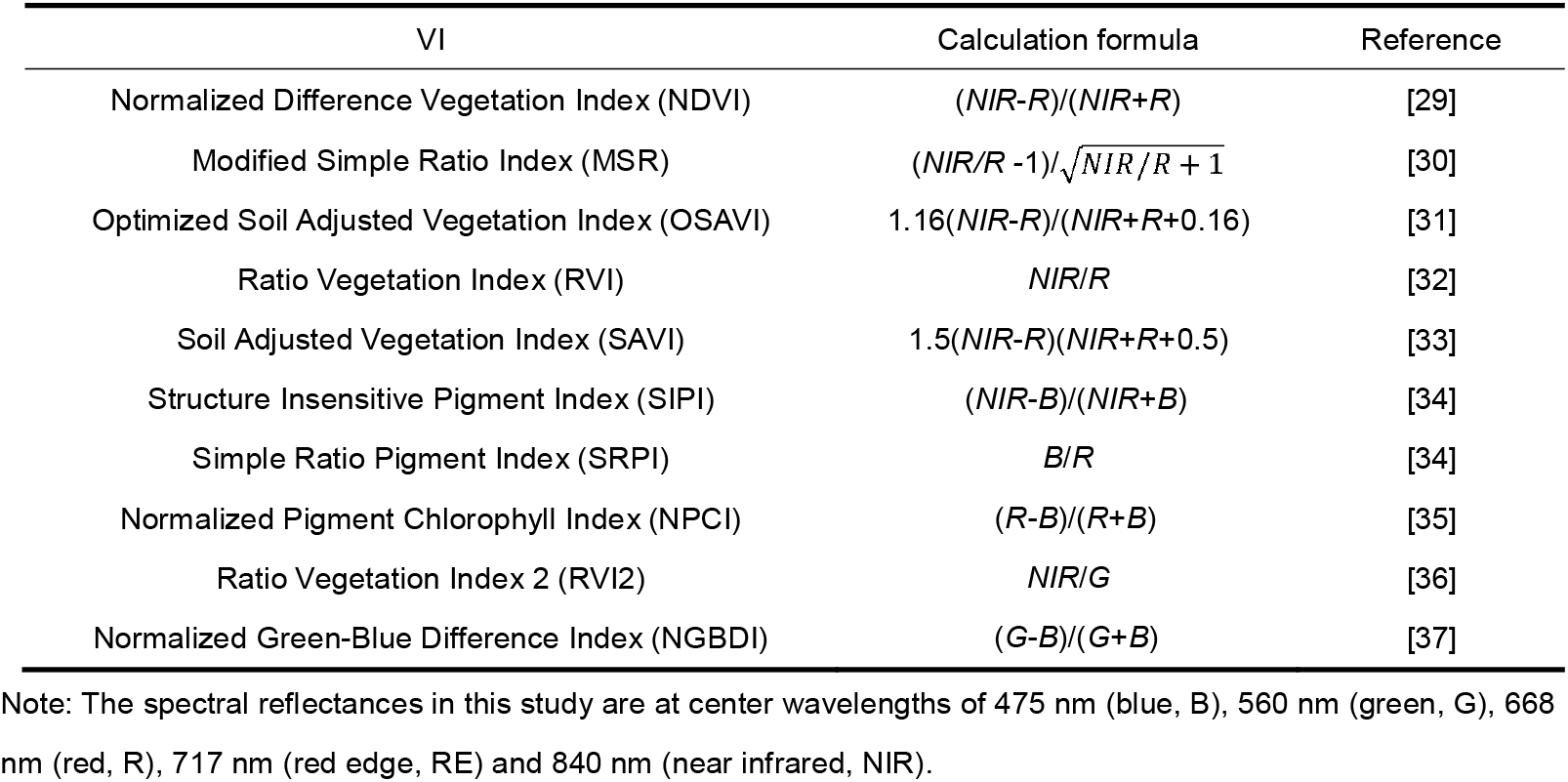
Selected VIs and their calculation formulas.

Grey relational analysis (GRA) is based on the development trend, and its requirements on the sample quantity and sample distribution are not strict. This process’s basic idea is to determine the primary and secondary relationships between various factors by calculating the grey relational degree (GRD) to determine the factor with the greatest influence. The higher the GRD of any two factors is, the more consistent the change between those two factors.

Let the reference sequence be *x*_o_ = {*x*_o_(*k*),*k* = 1,2,…, *n*} and the comparison sequence be *x_i_* = {*x_i_* (*k*),*k* = 1,2, …, *n*}. The GRD between *X_0_* and *X_i_* is calculated by Equations 1 and 2.

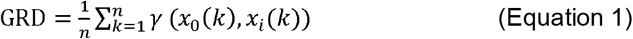

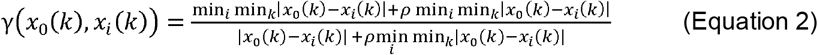

where *ρ* is the grey correlation coefficient, which ranges from 0-1; here, we took 0.5 as the value 38.

All the samples’ N contents were used as the reference sequence, and all the selected VIs were used as the comparison sequences. The GRDs between each VI and the N content are listed in Table 3.

**Table 3.** GRA result between each VI and N content.

| VI | GRD | Rank |
| --- | --- | --- |
| NGBDI | 0.8891 | 1 |
| SRPI | 0.8053 | 2 |
| NPCI | 0.6375 | 3 |
| SIPI | 0.6180 | 4 |
| RVI | 0.6049 | 5 |
| NDVI | 0.5798 | 6 |
| RVI2 | 0.5797 | 7 |
| MSR | 0.5503 | 8 |
| OSAVI | 0.5400 | 9 |
| SAVI | 0.5154 | 10 |

The NGBDI had the strongest relation with the N content. The first six VIs, namely, the NGBDI, SRPI, NPCI, SIPI, RVI, and NDVI, were selected for N prediction modeling (Table 3). Different combinations of VIs were used to build the single VI prediction models, multiple VI prediction models, hybrid prediction models with raw band reflectance data. All the results are listed in Tables 4 and 5. It is shown that

1. The R values of the models built from the VIs were approximately 0.3-0.5, showing poor accuracy, but the models combining the VIs with raw band reflectance data had obviously higher R values (approximately 0.5-0.8) and lower RMSE values (approximately 0.1-0.2) than the models built with only the VIs.
2. The model with the best performance was built with the PLS algorithm and the raw band data, NGBDI, SRPI, and NPCI. This model had the highest R value (0.7693), obviously lower RMSE (0.1109) and SEP (0.1147) values, and the most balanced accuracy between the training set and validation set compared to the other algorithms. This result is similar to many other studies on other crops based on multispectral images, such as winter wheat N prediction (R^2^=0.53, RMSE=0.28) by Lu et al. [39], grapevine leaf N prediction (R^2^=0.56, RMSE=0.23) by Moghimi et al. [40].
3. For every input variable set, the PLS model had the most balanced result compared to the BPNN and ELM models, as shown in Table 1. PLS was also demonstrated as the optimal algorithm because of its superior performance. PLS was used to predict citrus canopy N and reached a prediction accuracy (R value) of 0.6398 [41]. PLS combined with GRA was also used to estimate the accuracy of water content in winter wheat leaves, and the model effect was better than stepwise regression [42].

**Table 4.**
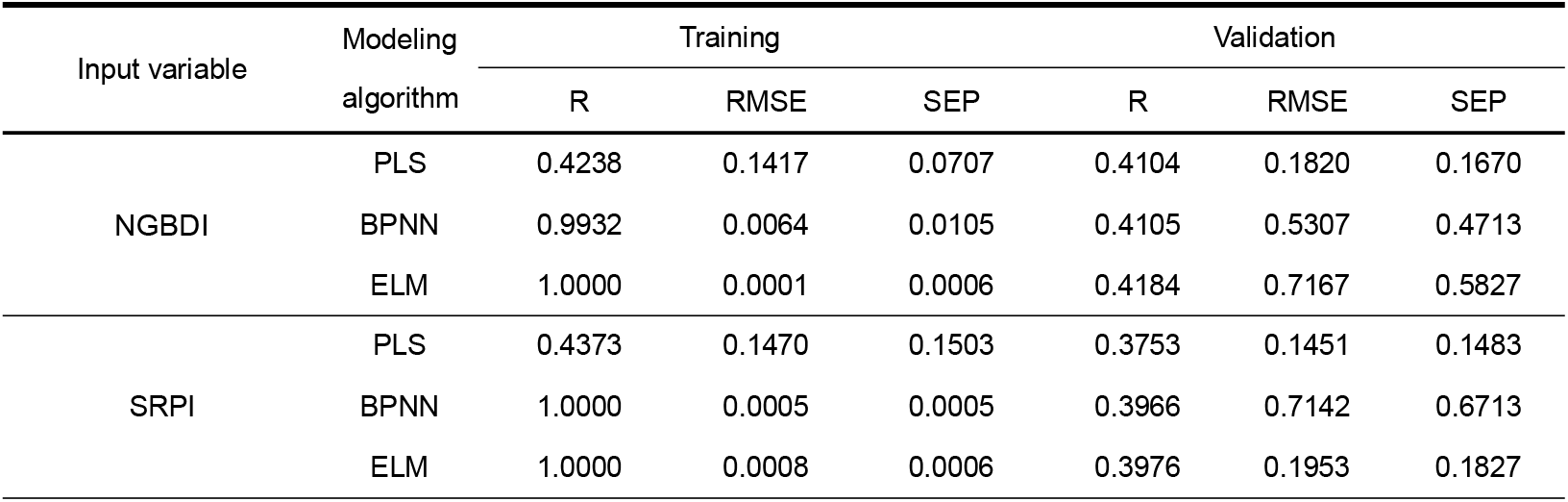
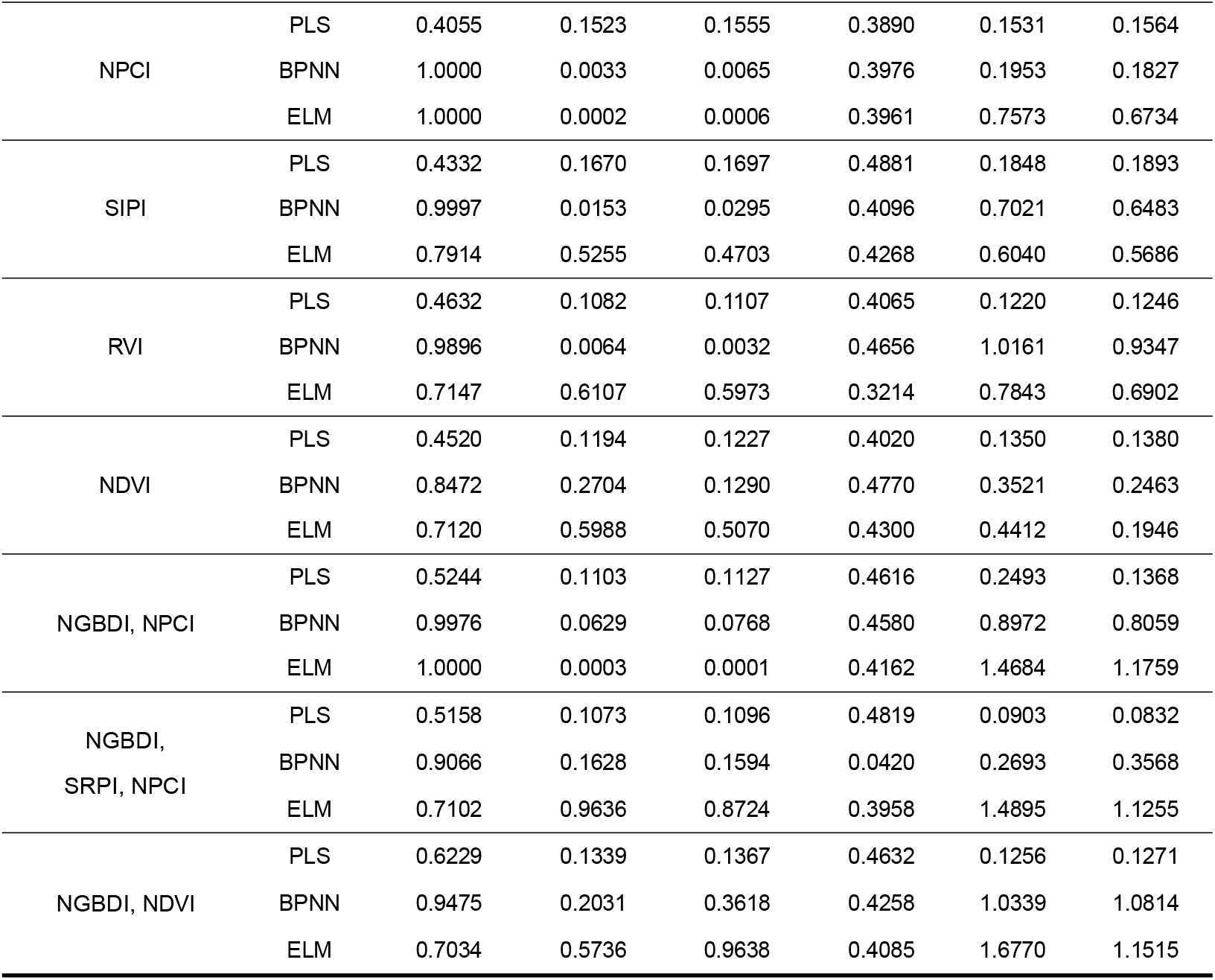
The modeling results from the VIs by using different algorithms.

**Table 5.**
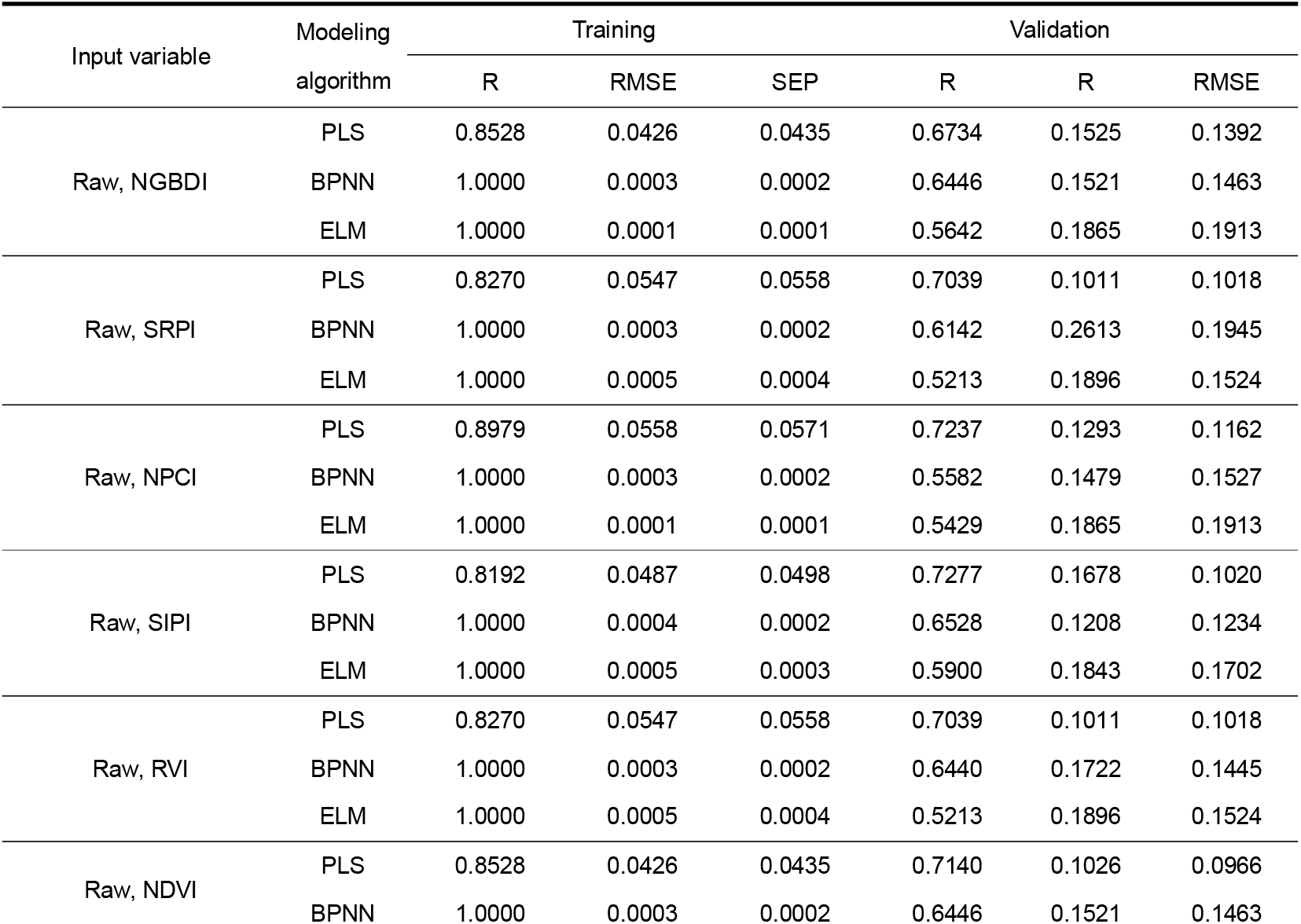
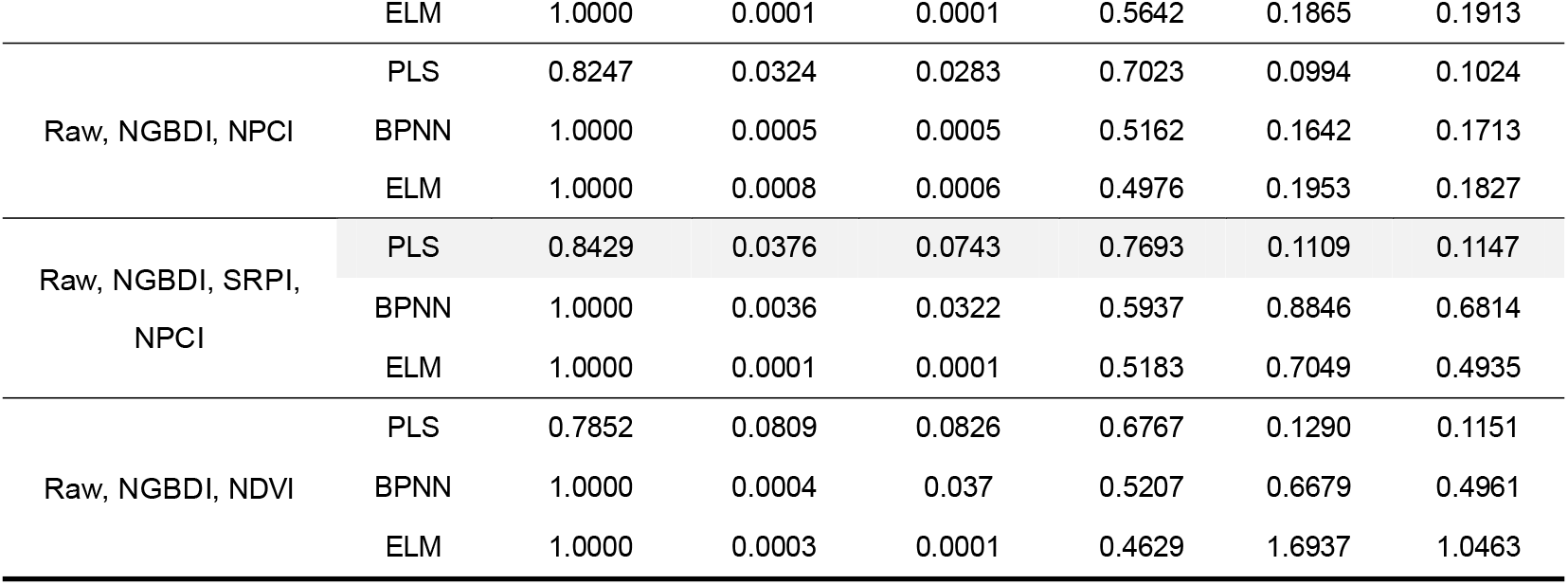
The modeling results from both the raw band reflectance and VIs by using different algorithms.

### 2.2 Irrigation level recognition based on multispectral images

Water is vital for sugarcane plants; water deficiency can cause obvious yield reduction [43]. In this research, the relations between spectral features and irrigation amounts were analyzed as well. First, scatter plots of all the band reflectances and VIs at different irrigation levels were generated. The red band, NDVI, RVI, MSR, SIPI, and RVI2 showed obvious differences under the different irrigation treatments, regardless of the amount of fertilizer applied, as shown in Fig 4. The red band shows a negative correlation with irrigation, while the other five VIs show a positive correlation. The average values of the same irrigation for all the spectral bands and VIs, as well as the correlations between average values and irrigation amounts, were also calculated (Table 6).

**Table 6.**
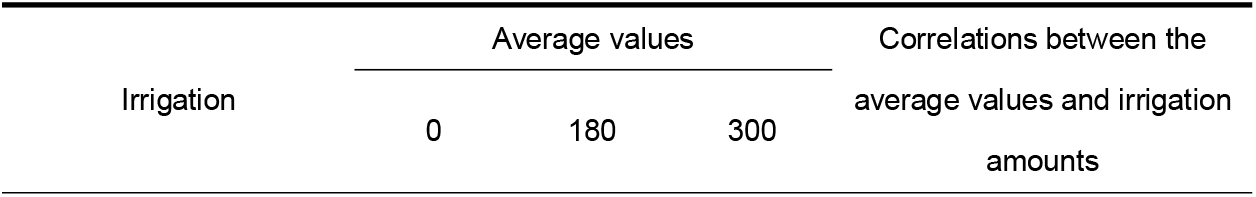
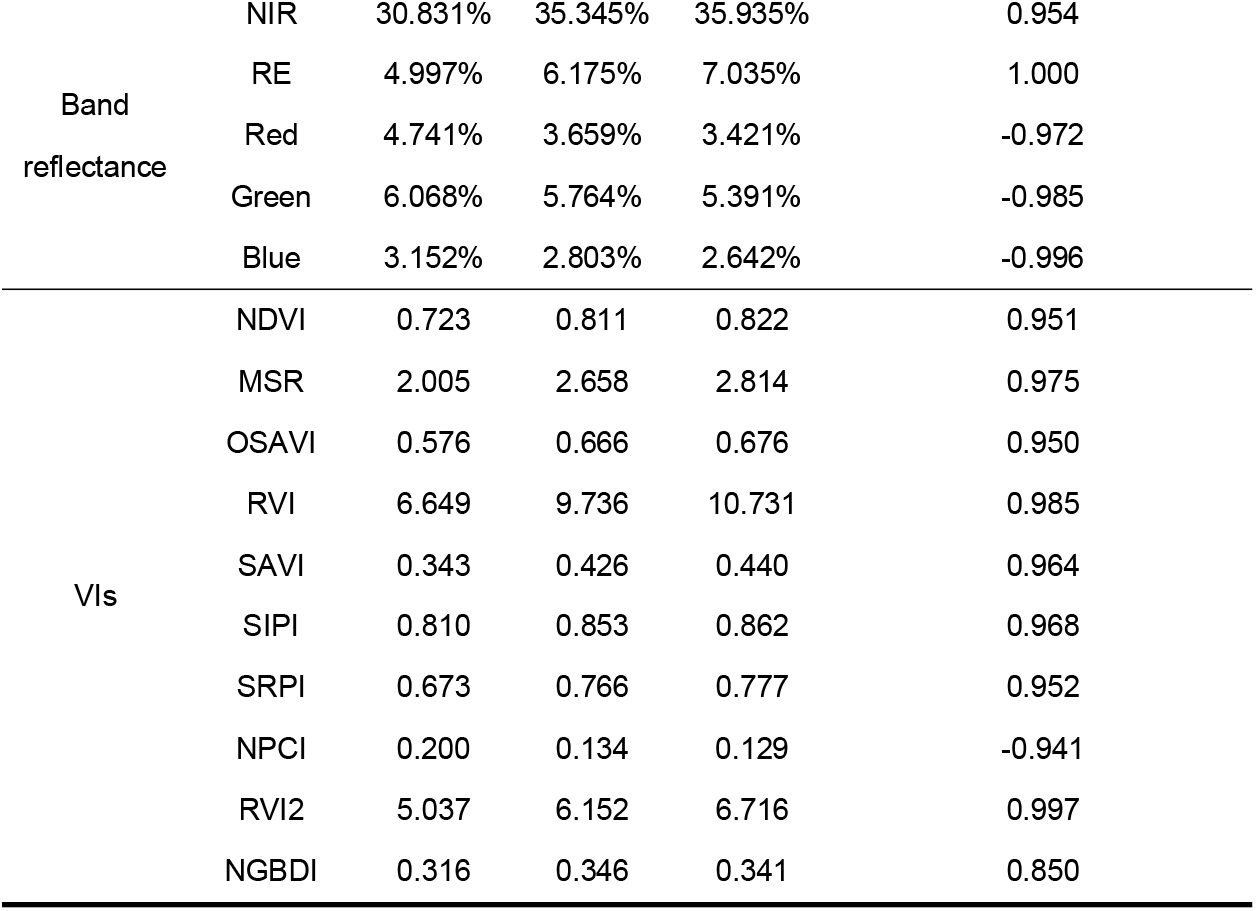
Average spectral values and their correlations with the irrigation amounts.

| Irrigation | Average values |  |  | Correlations between the average values and irrigation amounts |
| --- | --- | --- | --- | --- |
|  | 0 | 180 | 300 |  |

|  |  |  |  |  |  |
| --- | --- | --- | --- | --- | --- |
| Band<br>reflectance | NIR | 30.831% | 35.345% | 35.935% | 0.954 |
|  | RE | 4.997% | 6.175% | 7.035% | 1.000 |
|  | Red | 4.741% | 3.659% | 3.421% | -0.972 |
|  | Green | 6.068% | 5.764% | 5.391% | -0.985 |
|  | Blue | 3.152% | 2.803% | 2.642% | -0.996 |
| VIs | NDVI | 0.723 | 0.811 | 0.822 | 0.951 |
|  | MSR | 2.005 | 2.658 | 2.814 | 0.975 |
|  | OSAVI | 0.576 | 0.666 | 0.676 | 0.950 |
|  | RVI | 6.649 | 9.736 | 10.731 | 0.985 |
|  | SAVI | 0.343 | 0.426 | 0.440 | 0.964 |
|  | SIPI | 0.810 | 0.853 | 0.862 | 0.968 |
|  | SRPI | 0.673 | 0.766 | 0.777 | 0.952 |
|  | NPCI | 0.200 | 0.134 | 0.129 | -0.941 |
|  | RVI2 | 5.037 | 6.152 | 6.716 | 0.997 |
|  | NGBDI | 0.316 | 0.346 | 0.341 | 0.850 |

**Fig 4.**
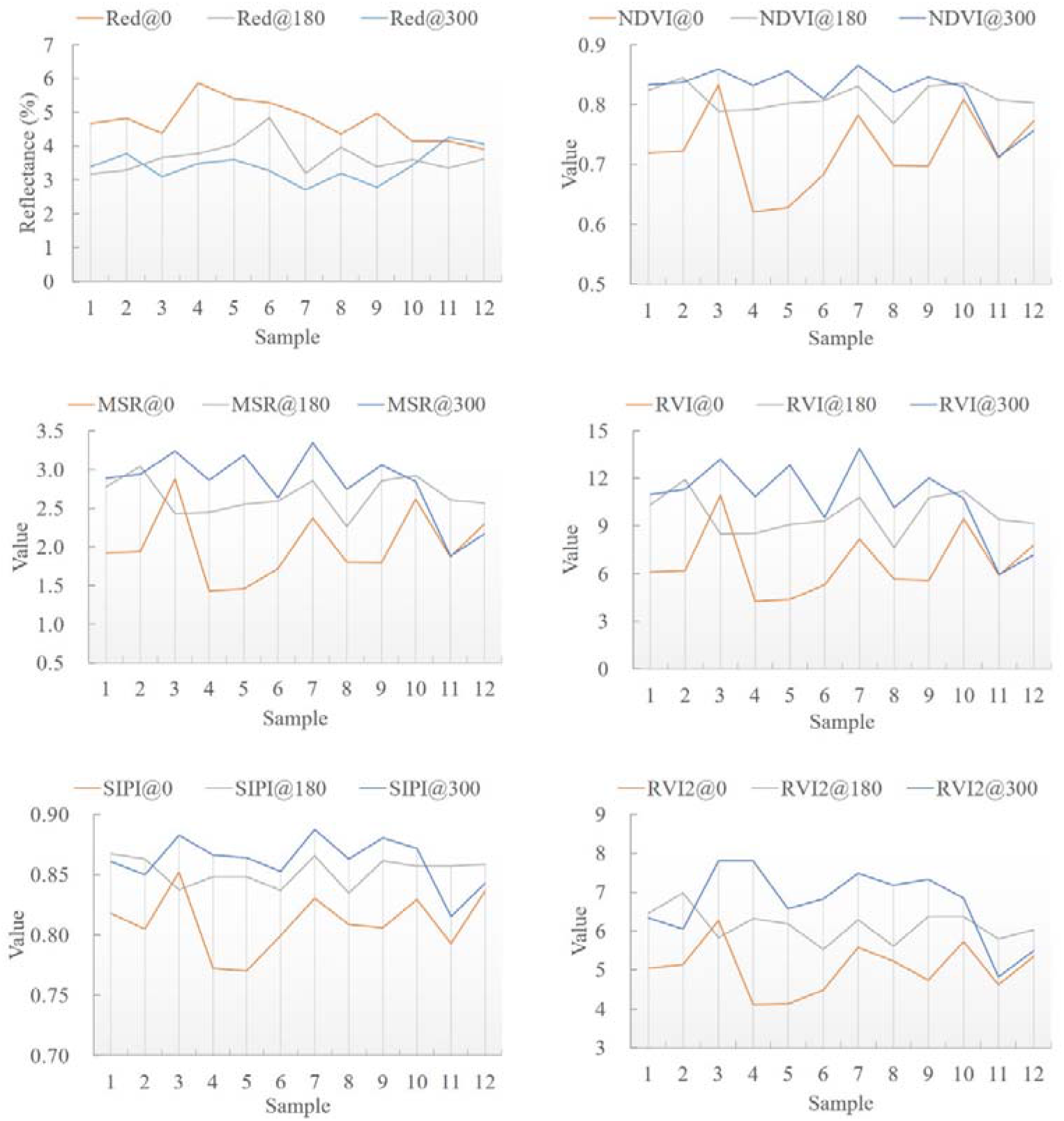
Scatter plots of the red band and five VIs according to the different irrigation levels. 0 represents rain with no drip irrigation; 180 represents a drip irrigation amount of 180 m^3^/ha (W0.6); 300 represents a drip irrigation amount of 300 m^3^/ha (W1.0).

For the average reflectance data of spectral bands, the mean reflectance of visible bands (red, green, and blue) decreased as the irrigation amount increased, showing strong negative correlations, while the other two bands (near infrared (NIR) and red edge (RE)) showed a strong negative trend (Table 6). Normally, green plants have lower reflectance in the visible range and higher reflectance in the NIR range as they are more vigorous. This result indicates that a proper increase in irrigation amount could make sugarcane healthier. For the average VIs, most VIs had strong correlations with the irrigation amounts demonstrating high possibility for irrigation level prediction.

The obvious difference in sample scale shown in Fig 4 indicate that the red band, NDVI, RVI, MSR, SIPI, and RVI2 could have great potential to classify different irrigation levels. An SVM was used to implement this classification. Before building the classification model, all six variables were normalized and then selected as the model inputs. The classification results are listed in a confusion matrix, as shown in Table 7. Each row represents the predicted irrigation levels; the user’s accuracy (UA) is the ratio of correctly classified irrigation levels to the total number of classified irrigation levels for that class. Each column represents the actual irrigation levels; the producer’s accuracy (PA) is the ratio of correctly classified irrigation levels to the total number of actual irrigation levels for that class. The overall classification accuracy was 77.8%; most of the misclassified samples were misclassified as adjacent irrigation levels, and only one sample was misclassified as a very different level (a sample from Irrigation_300 was misclassified as Irrigation_0), showing the great potential of irrigation level recognition with multispectral images.

**Table 7.** Confusion matrix of the irrigation level classification.

| Classification result \ Ground truth |  |  |  |  |  |
| --- | --- | --- | --- | --- | --- |
|  | Irrigation_0 | Irrigation_180 | Irrigation_300 | Total | UA |
| Irrigation_0 | 9 | 0 | 1 | 10 | 90% |
| Irrigation_180 | 3 | 11 | 3 | 17 | 64.7% |
| Irrigation_300 | 0 | 1 | 8 | 9 | 88.9% |
| Total | 12 | 12 | 12 | 36 |  |
| PA | 75% | 91.7% | 66.7% |  | 77.8% |

## 3 Conclusions

In this paper, a low-altitude UAV was used to collect multispectral images of sugarcane fields. Three machine learning models with band reflectance and multiple VIs as input variables were used to build N content prediction models, and their prediction performances were compared. In addition, an irrigation level recognition algorithm based on multispectral images was also proposed. The main conclusions were as follows:

1. The combination of the VI and band reflectance had better prediction accuracy than the VI or band reflectance alone.
2. The PLS algorithm showed the best performance for sugarcane N prediction. The PLS algorithm built the best model from the raw band data, NGBDI, SRPI, and NPCI, which had the highest R value of 0.7693, an obvious lower RMSE value of 0.1109, and most balanced accuracy between the training set and validation sets.
3. Sugarcane fields with different irrigation levels had an obvious difference in the spectral information of their multispectral images. An SVM classification model was built to classify the irrigation levels, and the classification accuracy reached 77.8%.

Multispectral images acquired by low-altitude UAVs could provide effective information for N content prediction and water irrigation level recognition. Further research should be performed based on multitemporal spectral images and more comprehensive ground investigation.

## Acknowledgments

This work was supported by the Science and Technology Major Project of Guangxi, China (Gui Ke 2018-266-Z01), the National Natural Science Foundation of China (31760342, 31760603), and the Science and Technology Major Project of Guangxi, China (Gui Ke AA18118037).

## Author Contributions

Conceptualization: Xiuhua Li

Investigation: Yuxuan Ba, Shimin Zhang, Mengling Nong, Ce Wang

Methodology: Yuxuan Ba, Xiuhua Li

Writing – Original Draft Preparation: Xiuhua Li, Yuxuan Ba

Supervision, Writing – Review & Editing: Xiuhua Li, Muqing Zhang

## Notes

### Competing Interest Statement

The authors have declared no competing interest.

### Summary of Updates

The paper was edited for grammar, phrasing, and punctuation. In addition, many edits were made to further improve the flow and readability of the text.

